# Intra-lineage microevolution of *Wolbachia* leads to the emergence of new cytoplasmic incompatibility patterns

**DOI:** 10.1101/2023.06.08.544196

**Authors:** Alice Namias, Annais Ngaku, Patrick Makoundou, Sandra Unal, Mathieu Sicard, Mylène Weill

**Affiliations:** ISEM, Université de Montpellier, CNRS, IRD, EPHE, Montpellier, France

**Author notes:** Corresponding authors, (AN), (MW).

**Keywords:** *Wolbachia*, *Culex pipiens*, cytoplasmic incompatibility, microevolution, recombination

## Abstract

Mosquitoes of the *Culex pipiens* complex are worldwide vectors of arbovirus, filarial nematode, and avian malaria agents. In these hosts, the endosymbiotic bacteria *Wolbachia* induce cytoplasmic incompatibility (CI), that is, reduced embryo viability in so-called incompatible crosses. *Wolbachia* infecting *Culex pipiens* (*w*Pip) cause CI patterns of unparalleled complexity, associated with the amplification and diversification of *cidA* and *cidB* genes, with up to six different gene copies described in a single *w*Pip genome. By repeating crosses between *Culex* isofemale lines over 17 years, we documented the emergence of a new compatibility type. Using a new sequencing method adapted to multigene families to acquire *cid* genes, we showed that some *w*Pip genomes lost specific *cidA* gene copies, thus giving rise to several sub-lineages segregating in the same cage. By linking phenotypic changes to their underlying genotypic bases, we showed that gene copies that are key for CI phenotypes originated from recombinations, not point mutations. We revealed how new CI patterns could emerge as part of a two-step process: first, local changes take place in the CI repertoires while maintaining compatibility with the surrounding mosquitoes, and then migration and secondary contact occur with the incompatible lines.

## Introduction

*Wolbachia* are maternally transmitted endosymbiotic bacteria that infect up to 50% of arthropod species [1–3]. These bacteria are well known for their wide range of reproductive manipulations in arthropods. Their most common manipulation is cytoplasmic incompatibility (CI), that is, a reduction in embryo hatching rates in crosses between infected males and uninfected females, which is the simplest form of CI. CI results from a *Wolbachia*-induced *modification* that perturbs the first division of embryos, which infected females are able to rescue [4,5]. CI is thus formalized in a *modification-rescue* framework.

CI can also occur between two infected individuals if they are infected with genetically different and incompatible *Wolbachia* strains. In *Culex pipiens* mosquitoes, in which *Wolbachia* infection is fixed [6], the *w*Pip strains are responsible for a unique (to date) complexity of CI patterns based on multiple uni- and bidirectional incompatibilities [7–10]. An initial analysis of *w*Pip genetic diversity in *Culex* identified five phylogenetic groups named *w*Pip-I to *w*Pip-V [11]. Comparing *w*Pip phylogeny with crossing results revealed that these groups were good predictors of compatibility outcomes [9]: while both compatible and incompatible crosses were reported in crosses among *w*Pip groups, intra-group crosses were, with rare exceptions, compatible.

In 2013, putative CI genes were identified using a combination of proteomic and genetic approaches [12]. Then, in 2017, two functional studies confirmed that these genes were key for CI, naming them cif for “CI factors” [13,14]. The *cif* genes were described in *cifA*-*cifB* tandems, playing a central role in CI, although the precise molecular mechanisms of CI are still to be determined. Two main alternative models are still being debated: a host-*modification* (HM) model in which *Wolbachia* toxicity is due to the *modification*s of the host, with rescue occurring through a reversal of these *modification*s; and a toxin-antidote (TA) model in which the sperm carries a *Wolbachia* factor that is toxic unless an appropriate *Wolbachia* antidote is expressed in the egg, thus binding and neutralizing the toxin [15–18]. In *Culex*, the complexity of the crossing patterns makes the TA model far more likely: (in)compatibility would thus be based on different toxin-antidote binding abilities [16,19–21]. Recent publications agree with *cifA* being the rescue (resc, or antidote) gene and *cifB* being a major *modification* (mod, or toxin) gene [13,18,22]. Sequencing of *Wolbachia* genomes from different host species revealed several different pairs of *cif* genes, with distinct functional domains that categorized them into five clades [23–25]. All *w*Pip genomes sequenced so far have *cif* genes from two of these clades: clade I cif, in which the *cifB* gene has a deubiquitinase domain (the tandem is thus called *cid*, with d denoting the deubiquitinase domain), and clade IV cif, in which *cifB* bears a nuclease domain (thus called cin) [13,14]. Here, we use this functional-based nomenclature [19].

Before the discovery of the *cif* genes, models based on *Culex*-crossing experiments predicted that several factors or pairs of *mod/resc* genes were required to encode the complex crossing patterns induced by *w*Pip [9,26]. Studies on *cif* (*cid* and *cin*) genes present in *w*Pip genomes highlighted that cin genes were always monomorphic, whereas *cid* genes were amplified and diversified in all sequenced *w*Pip genomes [27], with up to six different copies of each gene in a single *Wolbachia w*Pip genome. The different *cidA/cidB* copies in a given *w*Pip genome are known as “variants” and the full set of all copies constitutes the “*cid* repertoire.” In *w*Pip genomes, the polymorphism of both *cidA* and *cidB* genes are located in two specific regions, which were named “upstream” and “downstream” regions and predicted to be involved in *cidA*-*cidB* interactions [27]. This has been confirmed by the recently obtained structure of a *cidA*-*cidB* cocrystal: out of the three interaction regions identified, two perfectly match the previously identified upstream and downstream regions for both *cidA* and *cidB* [20,21]. In addition to showing that *cidA* and *cidB* from *w*Pip bind together, this recent study also showed that the different *cidA*-*cidB* variants have different binding properties, thus lending further support to the toxin-antidote model for *w*Pip CI complexity [21].

Evolutionary changes of *Wolbachia* genomes on long time scales have been widely documented in different host species. The comparison of *cif* genes in *Wolbachia* strains from different arthropod hosts showed that they are quite divergent, and highly subject to lateral gene transfers [28,29] so that the congruence between *Wolbachia* and *cif* phylogenies is totally disrupted [23,24]. Such lateral transfers may be phage-linked, as *cif* genes are located in WO prophage regions [14,23], with lateral transfers of genes in prophage WO regions being previously documented [30–33]. Transposon-dependent (and thus phage-independent) transfers of *cif* genes have also been described [28]. By contrast, although the ability of *Wolbachia* to rapidly adapt to new environmental conditions has been suggested [34], very few studies have explored the short-term evolution of *Wolbachia*. To our knowledge, the only studies linking rapid changes in phenotypes (in a few host generations) to underlying genomic variations were reported for the “Octomom region” in *w*Mel and *w*MelPop, showing that variations in the amplification level of this region were responsible for variations in both *Wolbachia* virulence and viral protection conferred by *w*Mel [35–37].

CI patterns in *Culex* were previously shown to change over a few host generations [38]. Yet these changes were described before the discovery of *cif* genes. Here, we observed a change in the CI pattern between two isofemale lines kept in our laboratory since 2005: Slab (*w*Pip III) and Istanbul (Ist, *w*Pip IV) [10]. While crosses between Slab females and Ist males were compatible from 2005 to 2017, we observed fully incompatible crosses for the first time in 2021. This shift in CI patterns presented an opportunity to study the underlying genetic basis of CI evolution in laboratory-controlled isofemale lines.

The emergence of a new CI phenotype may have resulted from (i) contamination, (ii) the acquisition of a new toxin in some Ist *Wolbachia*, or (iii) the loss of antidotes in some Slab *Wolbachia*. We took advantage of a recent methodological development that enables the rapid and extensive acquisition of *cidA* and *cidB* repertoires using Nanopore Technologies sequencing [39] to address these three hypotheses. We were able to rule out the contamination hypothesis and show that no less than three distinct *Wolbachia* sublineages with different *cidA* repertoires now coexist in the Slab isofemale line (but do not appear to coinfect the same individuals). We also found that the loss of *cidA* variants (i.e., antidotes) in Slab perfectly matches variations in their rescuing ability. The *cidA* variant whose presence/absence best matches the CI variations has original aminoa*cid* combinations at its interaction interface with *cidB*, probably resulting from a recombination.

## Results

### From CI phenotype shift to underlying genomic basis

In 2021, crosses between Slab females and Ist males highlighted that some of the Slab females were fully incompatible with Ist males for the first time.

### At least three *cidA* repertoires coexist in the Slab isofemale line

To decipher the genomic changes that led to this recent CI shift, we sequenced the *cidA* and *cidB* repertoires present in *Wolbachia* infecting four individuals from the Ist isofemale line and eight females from the Slab isofemale line (four compatible and four incompatible) using Nanopore Technologies sequencing of *cidA* and *cidB* polymerase chain reaction (PCR) products [39]. While *Wolbachia* present in all Ist individuals had the same *cid* repertoires (S1 Table), we found polymorphism in *cid* repertoires of Slab individuals: they all had the same *cidB* (i.e., toxin) diversity, composed of *cidB-III-ae3* and *cidB-III-ag1* variants, whereas *cidA* (i.e., antidotes) repertoires varied among individuals. *Wolbachia* present in compatible and incompatible Slab females all exhibited the variants *cidA-III-alpha(5)-25* and *cidA-III-gamma(3)-12*, although additional variants were found in compatible females: *cidA-III-beta(2)-25*, either alone or accompanied by *cidA-III-beta(2)-16*. The observed differences in *cidA* repertoires can be summed up by the presence/absence of two specific *cidA* regions: the upstream region *cidA-III-beta(2)* and the downstream region *cidA-III-16* (S1 Fig).

### *cidA* repertoire variation fully explains compatibility polymorphism

To determine if compatibility variations were linked to these specific *cidA* repertoire variations, we performed three replicates of the [Slab x Ist] cross. We isolated egg rafts from a total of 102 Slab females from the same isofemale line (28, 34, and 40 females for replicates 1, 2, and 3, respectively) and studied their hatching rates (HRs, i.e., number of hatched larvae over the number of eggs laid). Eight females produced unfertilized rafts and were thus excluded from further analyses. For the remaining 94 females, HRs ranged from 0% (fully incompatible raft) to 96.4%, thus highlighting the strong differences in the rescuing abilities of females. Of these 94 females, 25 (i.e., 27%) were fully incompatible with Ist males.

Since repertoires differed in the presence/absence of two specific regions of *cidA* repertoires, we designed primers and specifically screened the 94 females (S2 Table). The PCR assay confirmed that three distinct *Wolbachia cidA* repertoires coexisted in the Slab isofemale line: (i) individuals hosting *Wolbachia* with both the upstream *beta(2)* and downstream *16* regions, named (β+,16+); (ii) individuals hosting *Wolbachia* with only the upstream region *beta(2)* but not the downstream *16* region, named (β+,16-); and (iii) individuals hosting *Wolbachia* genomes with neither of these regions, named (β-,16-) (S2 Table). For the sake of simplicity, we metonymically qualified mosquitoes based on their *cidA* repertoires (β+,16+), (β+,16-), or (β-,16-). Of the 94 females analyzed, 61 were (β+,16+), 8 (β+,16-), and 25 (β-,16-) (Fig 1).

**Fig 1.**
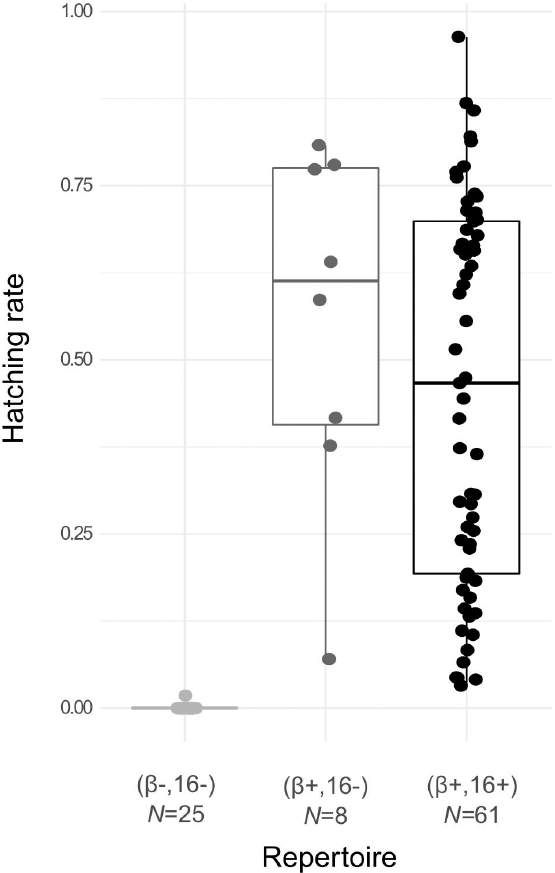
Intra-line polymorphism in *cidA* repertoires matches the compatibility patterns. (β-,16-) females have neither *cidA-III-beta(2)* nor *cidA-III-16* regions; (β+,16+) females have both regions, and (β+,16-) females have the *cidA-III-beta(2)* region but not the *cidA-III-16*. The presence or absence of *cidA-III-beta(2)* and *cidA-III-16* regions was determined by specific qPCRs. *N* = Number of individuals.

Egg rafts produced by (β-,16-) females had a null hatching rate, with a single exception, in which only one larva hatched, resulting in a HR of 1.8%. Fully incompatible crosses were only observed with (β-,16-) mothers, as all (β+,16+) and (β+,16-) mothers were compatible with Ist males (to varying extents) (Fig 1).

### *cidA* repertoire variations do not explain HR variability in compatible crosses

While it was long believed that the presence of CI in *Culex* was a “yes or no” phenomenon, a recent study explored HR variations in compatible crosses with *w*Pip from different groups [40]. Examining the compatible crosses involving (β+,16+) or (β+,16-) Slab females and Ist males, we also found strong variations in HRs ranging from 3.2% to 96.4% (S2 Table). While infection levels (quantified by qPCR) in full individuals were variable, with 0.11 to 10.1 *Wolbachia* genomes per host genome and an average of 1.0 ± 0.2 (mean ± standard error), this variability did not correlate with HR variations (S2A Fig, Spearman correlation test, ρ = −0.09, *p* = 0.49, S2 Table). Similarly, HR variations were not correlated with the *cidA* copy numbers (total *cidA*, *cidA-III-beta*, or *cidA-III-16* copy numbers) per *w*Pip-Slab genome investigated in Slab females (Spearman correlation test, ρ = −0.13, *p* = 0.28; ρ = 0.01, *p* = 0.94; ρ = −0.17, *p* = 0.16 for the total *cidA*, *cidA-III-beta*, and *cidA-III-16*, respectively; S2B-S2D Figs, S2 Table).

### Recombination in *cid* variants leads to changes in *cidA*-*cidB* interaction zones

Recent works revealed that there were three interaction interfaces between *cidA* and *cidB* proteins, named interfaces I to III (Fig 2A, [20,21]). The upstream region *cidA-III-beta(2)*, present in both (β+,16+) and (β+,16-), seems key to enable Slab females to rescue Ist males. This region corresponds to the first interaction interface of the *cidA* protein (Fig 2A). Looking closer at the protein sequences at this interface, we found that *cidA-III-beta(2)* has an original combination of aminoa*cid*s resulting from a recombination of the *cidA*-alpha and *cidA*-gamma regions (Fig 2B). Building a phylogenetic network of all the described *cidA* upstream regions [22,27,40,41] confirmed that beta-like regions were rare (only found in the *w*Pip group IV and Slab to date) and that they came from a recombination of the *cidA-gamma*-like and *cidA-alpha*-like regions, with these last two regions being frequent and found in all *w*Pip groups surveyed to date (Fig 2C). We also showed that the *cidA-delta* region was recombinant.

**Fig 2.**
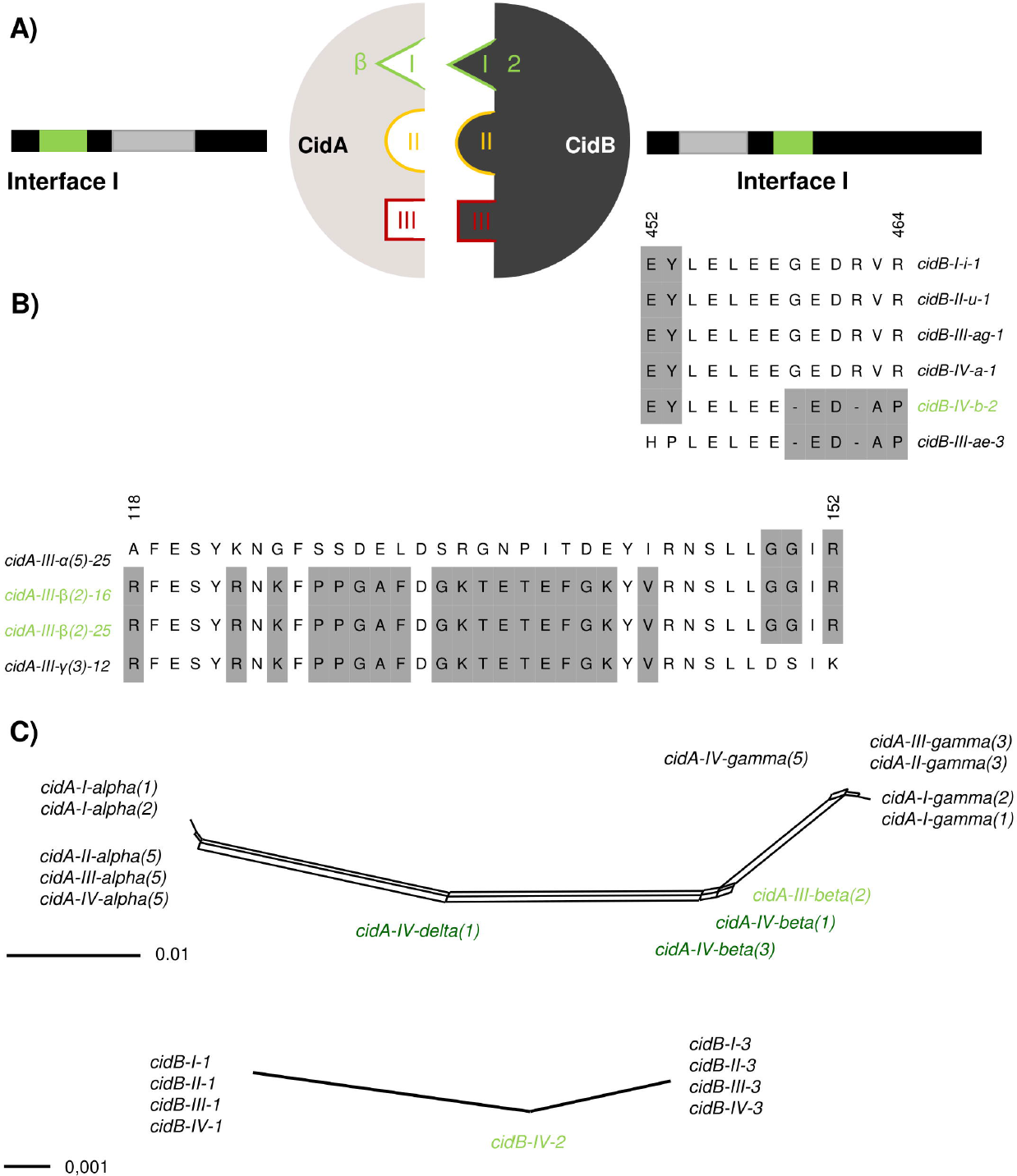
Matching recombination in the *cidA*-*cidB* interaction interface I is key for rescue. (A) *cidA* and *cidB* interact head-to-tail through three distinct interfaces, shown here in green, yellow, and red, respectively. The *cidA-III-beta(2)* and *cidB-IV-2* regions, which are key to the (in)compatibility between Slab and Ist isofemale lines, are both exposed in the interaction interface I. (B) Alignments of the protein sequences of the interaction interface I are shown for *cidA* and *cidB* (only sequence between the first and last variable aminoa*cid* belonging to interface I are represented; Wang & Hochstrasser, pers. com.). (C) Network phylogenetic analysis of the *cidA* upstream and *cidB* downstream regions using the Neighbor-Net method on 35 *cidA* variants and 22 *cidB* variants taken from this study and [27,40,41]. For both regions, there are two major groups of variants (alpha/gamma for *cidA* and 1/3 for *cidB*), with several intermediate recombinant variants (in green). Variants found in the present study are shown in light green. Each edge (or set of parallel edges) corresponds to a split in the dataset with a length equal to the weight of the split.

The *cidB*-IV-2 downstream region, present in *w*Pip-Ist, had previously been associated with incompatibility when crossed with a wide range of females [22]. This region interacts with *cidA-beta* at the interface I (Fig 2A) and also displays a pattern that could be explained by a recombination (between *cidB*-1 and *cidB*-3 downstream regions) (Fig 2B). When looking at wider phylogenetic scales based on all the previously published sequences, we found an even clearer pattern with only two *cidB*-1 and *cidB*-3 downstream regions, identical among *w*Pip groups, with *cidB*-IV-2 being the only intermediate recombinant region (Fig 2C).

### Evolution of *cid* repertoires

#### Different *cidA* repertoires segregate into sublines

Since a new CI pattern emerged between 2017 and 2021, a closer look at the transmission and variation of Slab individuals’ repertoires was required to better understand the precise nature of this evolution. For each of the three *cidA* repertoires found in the original Slab isofemale line, we established two isofemale sublines. We sequenced their *cidA* and *cidB* repertoires and confirmed that they were identical to those previously found for (β-,16-), (β+,16-), and (β+,16+) (S1 Fig). We confirmed that the females of these new isofemale sublines had the same CI phenotype as their respective founding females when crossed with Ist males (Table S3). We also verified that they were compatible with each other in both directions, showing that *cidA* repertoire polymorphism among sublines did not create incompatibilities (Table S4).

To test whether changes in repertoires could occur at the scale of a few host generations, we genotyped *Wolbachia* from 20 to 45 individuals of each Slab subline after about 15 generations for both *cidA* markers and found that no change had occurred since the establishment of the isofemale sublines.

We also genotyped individuals from the Slab isofemale line kept in liquid nitrogen since 2005, that is, shortly after the establishment of the isofemale line in our insectaries. Out of 30 tested mosquitoes, there were 26 (β+,16-) and 4 (β+,16+). None of them was (β-,16-), which is in accordance with the hypothesis that this genotype, leading to a new CI pattern, appeared afterwards.

#### No sign of stable coinfection

Using qPCR on Slab females, we found that total *cidA* copy numbers were variable, ranging from 2.2 to 6.2 copies per *w*Pip genome (quantifications normalized on the single copy *Wolbachia* gene wsp; S3A Fig). However, the distribution was bimodal with two peaks around three and five *cidA* copies (S3D Fig). The *cidA* copy number was homogeneous for females infected with a given *cidA* repertoire, with around five *cidA* copies for all (β+,16+) and (β+,16-) individuals and around three *cidA* copies for all (β-,16-) individuals (S3A Fig). We also quantified the number of *cidB* copies in eight to ten females of each repertoire and found that they had equal copy numbers of *cidB* and *cidA*: around five copies in both (β+,16+) and (β+,16-) individuals and around three *cidB* copies in (β-,16-) individuals (S4 Fig). These results show that while Slab *cidB* repertoires do not differ between lines in terms of the identity of variants, they differ in terms of the *cidB* copy numbers.

The variability in these qPCR results can be explained by the intrinsic variability of qPCR. The centered data distributions in S3 and S4 Figs support the conclusion that most individuals (if not all) from all the (β+,16+), (β+,16-), and (β-,16-) sublines are not coinfected by *Wolbachia* with different *cid* repertoires.

#### Multiple copies of the same variant in the repertoires

qPCR results revealed that the total number of *cidA* copies was always higher than the number of distinct variants, showing that some variants must be present in multiple identical copies in a given repertoire. By combining the qPCR data and relative read coverage information from Nanopore sequencing (e.g., variant with two copies is covered twice as much as one with a single copy), we estimated the copy numbers of each variant in the different *cidA* repertoires of the Slab line. We found that (β+,16+) individuals, which have a total of five *cidA* copies, had two copies of *cidA-III-alpha(5)-25* and a single copy of each of the other three variants (*cidA-III-beta(2)-16*, *cidA-III-beta(2)-25*, and *cidA-III-gamma(3)-12*). (β+,16-) individuals also had five *cidA* copies, of which there were two copies of *cidA-III-alpha(5)-25*, a single copy of *cidA-III-beta(2)-25*, and two copies of *cidA-III-gamma(3)-12*. Lastly, (β-,16-) individuals, which had a total of three *cidA* copies, had two copies of *cidA-III-alpha(5)-25* and a single copy of *cidA-III-gamma(3)-12* (Fig 3).

**Fig 3.**
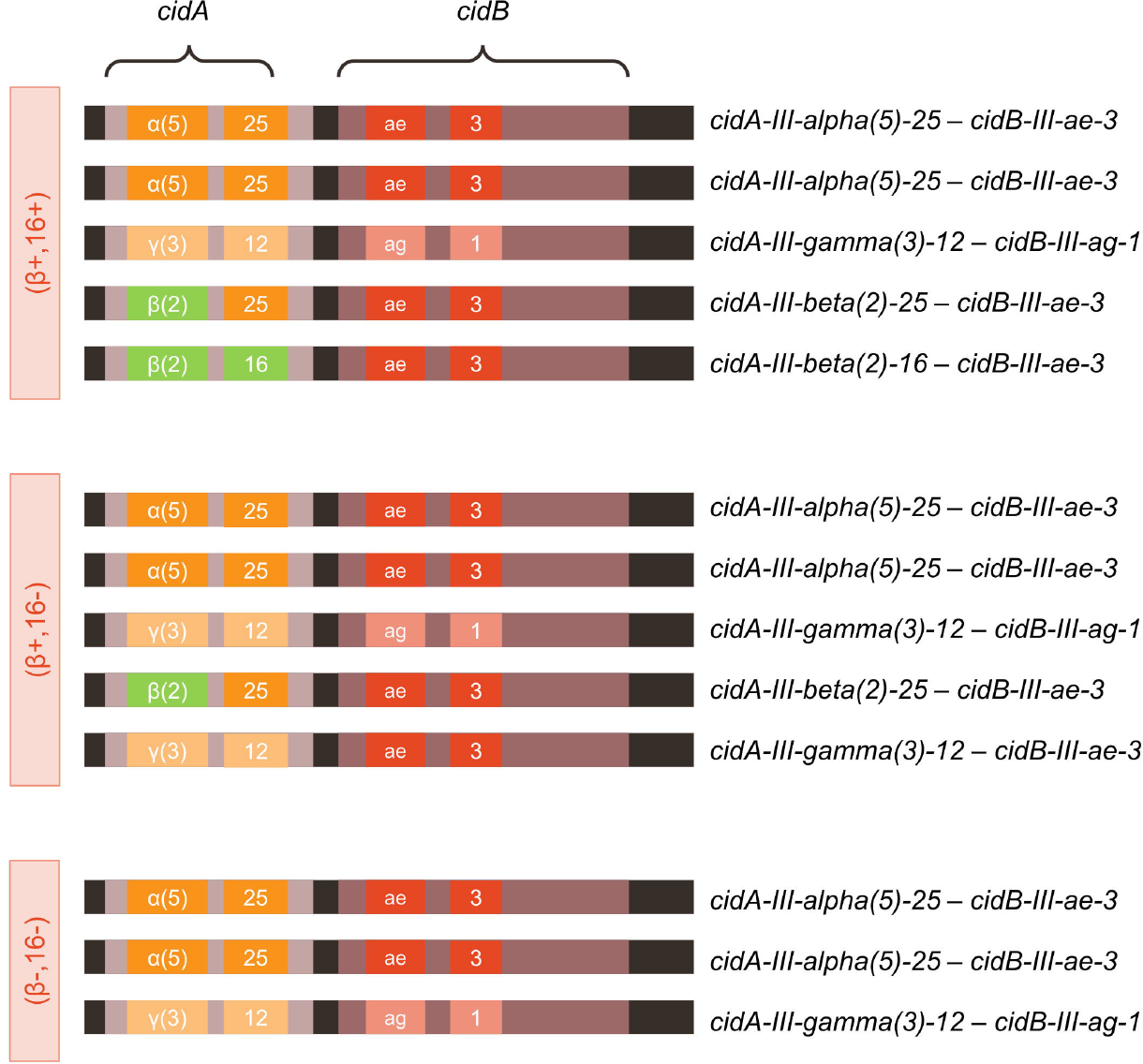
*cidA*B repertoires of *Wolbachia* infecting the three Slab sublines. The *cidA* gene is represented in light purple and *cidB* in dark purple. Upstream and downstream variable regions are shown in color. Regions in green are the key studied regions: *cidA-III-beta(2)* and *cidA-III-16*. The copy number of each tandem was deduced by combining the relative coverages in nanopore sequencing data with the qPCR data.

We confirmed the deduced copy numbers using qPCR specific to *cidA-III-beta(2)* and *cidA-III-16*: quantifying these two markers in 89 single Slab females, we confirmed that (β+,16+) repertoires had a single copy of *cidA-III-16*, while the other repertoires had none (S3B and S3E Figs). For *cidA-III-beta(2)*, we found that (β+,16+) individuals had two copies (corresponding to one copy of *cidA-III-beta(2)-16* and one copy of *cidA-III-beta(2)-25*), while (β+,16-) individuals had a single copy (unique copy of *cidA-III-beta(2)-25*). Finally, (β-,16-) individuals had none of these copies (S3C and S3F Figs).

#### *cidA* and *cidB* variants are always located within tandems

Previous investigations describing the *cid* repertoire of a given *Culex* line were performed by acquiring the sequences of *cidA* and *cidB* variants separately [22,27,39–41]. Here, we also amplified and Nanopore sequenced a region with both *cidA* and *cidB* variable regions in a single amplicon (S5 Fig). We could thus reconstruct the *cidA*-*cidB* tandems associated in the genome, resulting in the complete repertoires shown in Figure 3. To determine whether all *cid* variants were associated in tandems or whether they could stand alone in the genome, we compared their relative coverage when amplified and sequenced alone versus when amplified and sequenced in *cidA*-*cidB* tandem acquisitions. As these relative coverages were similar, we considered that all *cidA* and *cidB* variants were always located in tandems. Thus, (β+,16+) and (β+,16-) sublines have five *cidA*B tandems, while (β-,16-) lines have three *cidA*B tandems (Fig 3). Individuals of all sublines have *Wolbachia* with two copies of the *cidA-III-alpha(5)-25_cidB-III-ae3* tandem and one copy of the *cidA-III-gamma(3)-12_cidB-III-ag1* tandem. In addition, (β+,16+) and (β+,16-) individuals have one *cidA-III-beta(2)-25_cidB-III-ae3* tandem, although they differ in terms of their fifth tandem: *Wolbachia* in (β+,16+) individuals have a specific *cidA-III-beta(2)-16_cidB-III-ae3* tandem, while those in (β+,16-) individuals have a *cidA-III-gamma(3)-12_cidB-III-ae3* tandem instead (Fig 3). Our results show that the *cidB* variant “*cidB-III-ae3*” is associated in tandem with four different *cidA* variants (Fig 3).

## Discussion

The evolution of *Wolbachia* genomes has already been documented at large phylogenetic scales, with multiple horizontal transfers and gene losses (e.g., [31,32,37,42–45]). This is especially true for the *cif* gene family: numerous horizontal transfers have been described, resulting in a strong incongruence between *cif* gene phylogeny and *Wolbachia* phylogeny [23,24,28]. Although rapid shifts in CI patterns were previously described in *Culex* mosquitoes infected with *w*Pip [38], the *cif* genes had not been described at that time, thus preventing the study of the genetic basis of these phenotypic changes. Furthermore, we could not rule out the impact of contamination in such shifts. Here, we could link changes in CI patterns with their *Wolbachia* genomic basis for the first time. We detected a recent shift in CI patterns between Slab (*w*Pip-III) and Ist (*w*Pip-IV) lines: while they had been compatible for 12 years from 2005 to 2017, a quarter of the crosses were fully incompatible in 2021. We looked for underlying genomic variations in their *cid* repertoires. This shift in CI could have resulted from either a variation in the *cidB* (toxin) repertoires of Ist males or a variation in the *cidA* (antidote) repertoires of Slab females. We found that *cid* repertoires did not vary in Ist but that three different *cidA* repertoires coexisted in the same Slab isofemale line: two distinct repertoires with five *cidA* copies, labelled (β+,16+) and (β+,16-), which were able to rescue the *modification* induced by *w*Pip-Ist, and a repertoire with three *cidA* copies, named (β-,16-), which were unable to rescue these *modification*s. These results indicate that some *w*Pip genomes in the Slab isofemale line lost *cidA* variants. Indeed, the screening of mosquitoes from 2005 revealed that the (β-,16-) repertoire did not exist at that time, showing that some *cidA* variants were lost in some *w*Pip-Slab genomes between 2017, when all crosses were compatible, and 2021, when the shift in CI was detected.

By studying the genomic basis of phenotypic changes, we showed that *cid* gene repertoires truly evolved in *w*Pip over a period of 4 years. While previous studies showed that *cid* genes are highly variable within and between *w*Pip groups (with up to 30 distinct variants described to date for *cidA* and 20 for *cidB* [22,27,40,41]), we found that the new (β-,16-) repertoire is composed of *cidA* and *cidB* variants present in ancestral repertoires, which are associated in an identical *cidA*-*cidB* tandem architecture. We thus conclude that the shift in the CI pattern results from the evolution of the repertoire and not from any type of contamination.

### Rapid evolution of *cid* repertoires through gene losses

In 2017, approximately one quarter of females had a total of three *cidA*-*cidB* tandems, showing that two *cid* tandems had been lost. *cid* copy number distributions are bimodal, with two peaks at three and five copies and very rare with intermediate repertoires of four copies. These rare intermediates are likely due to the qPCR variability. This could mean either that (i) four-copy repertoires were a transient state that disappeared quickly due to lower fitness or drift or that (ii) four-copy repertoires did not exist, meaning that two tandems were lost simultaneously.

Here, we found that all the observed repertoires were mutually compatible. This means that mosquitoes infected by the putative four-copy repertoires would also have been compatible with all surrounding mosquitoes, making its quick disappearance unlikely. The most plausible hypothesis is that two tandems were lost simultaneously, making it likely that the two tandems were located next to each other.

Literature on CI evolution suggests that mod factors or toxins (cifB variants) could degenerate first, as their evolution is neutral for maternally transmitted *Wolbachia* [46,47]. Only once *cifB* variants have lost their function is it possible for the loss of *cifA* (resc factors or antidotes) [48]. In such a conceptual framework, losing *cifA* without losing *cifB* would lead to self-incompatibility, which is an immediate evolutionary dead-end. Independent from CI, the presence of *cidB* without an associated *cidA* might be toxic for the host, as shown recently in insect cells [4]. These predictions have been corroborated by a large-scale phylogenetic study of *cif* genes, indicating that *cifB* genes were often disrupted, while *cifA* were not [24]. However, at a microevolutionary scale, we observed that it was possible to qualitatively decrease the diversity of *cidA* variants without qualitatively changing the *cidB* variants: all (β+,16+), (β+,16-), and (β-,16-) repertoires described here share the same *cidB* variants (*cidB-III-ag1* and *cidB-III-ae3*), while they have four, three, and two distinct *cidA* variants, respectively (Fig 3). This apparent discrepancy between the predictions and our results may be explained by the fact that *w*Pip genomes contain several *cid* genes. Unlike the other systems studied to date where the loss of *cifA* can lead to a loss of the self-rescuing ability, *w*Pip has several *cidA* variants, which makes it possible to lose one variant while still maintaining compatibility with the surrounding mosquitoes. The loss of *cidA* variants required for intra-line compatibility may also occur, although the mosquitoes that lost these key antidotes would be quickly wiped out by selection, a hypothesis that has been corroborated by the mutual compatibility between all newly established isofemale lines, regardless of their *cid* repertoire.

Laboratory conditions could facilitate this evolution by qualitatively reducing the *cidA* repertoire. Indeed, in natural populations, multiple *cid* repertoires coexist (as shown by sequencing and revealed by numerous crosses [22,27,41,49]), meaning that individuals are exposed to a wide range of toxins. A greater diversity in the *cidA* repertoire enables the host to rescue a wider range of males, which is thus under positive selection. While *cidA* diversity is advantageous in natural populations, in a cage with an isofemale line, females are exposed to a restricted and fixed set of toxins. Loss of *cidA* variants, which are not strictly required for the rescue of the *cidB* in a given cage, is not counter-selected. Furthermore, as numerous gene copies can be costly (e.g., [50]), the loss of *cidA*-*cidB* tandems could confer a higher fitness to *Wolbachia* with lower numbers of copies and thus contribute to the fixation of new *Wolbachia* genomes in mosquitoes, which could explain the rapid *Wolbachia* turnover. It is conceivable that the three-copy repertoire of (β-,16-) *Wolbachia*, which contains two distinct variants of *cidA* and two distinct variants of *cidB*, is the minimum viable repertoire with only pairs of matching toxins and antidotes.

### No lasting coinfection involved in changes in CI patterns in *w*Pip

One plausible explanation for the observed shift in CI could be that Slab is coinfected by two strains of *Wolbachia* with different compatibility types and different *cid* repertoires, which are usually co-transmitted, and that some individuals recently lost one of these strains, as observed in *Aedes albopictus* [51,52]. Using common MLST markers, Baldo et al. initially found no variability in the *w*Pip strains [53], showing that if coinfection occurs, it should be between highly similar strains. The development of a *w*Pip-specific MLST led to the uncovering of *w*Pip polymorphism and the description of five monophyletic groups within *w*Pip [11]. However, while these markers highlighted *w*Pip polymorphism, no sign of coinfection was found, as every mosquito clearly hosted one of the five groups [11]. In this study focusing on *cid* genes, which are highly variable even within a *w*Pip group, we likewise found no clear signs of coinfections. We quantified several *cid* markers and analyzed the distribution of their values. In the case of coinfections with two *Wolbachia* strains with three and five *cid* copies, respectively, quantification outcomes would have varied depending on the proportion of each *Wolbachia* in each mosquito, leading to a unimodal distribution centered on an intermediate value. For the three distinct *cid* markers analyzed in 89 females, the quantifications displayed a bimodal distribution (S3D-F Figs), with a few intermediate values probably due to technical variations in the qPCR measures. While coinfection occurred at some point (all *Wolbachia* within one individual cannot be wiped out and instantly replaced by another *Wolbachia*), it had to be brief with no lasting coinfection in a single host individual. This could be explained by two alternative hypotheses: (i) an exclusion between the two distinct genotypes (three and five copies) for any non-CI-related reason, or (ii) a small effective population of *Wolbachia* (number of *Wolbachia* transmitted from mother to offspring) in *w*Pip.

### Variability in hatching rates in compatible crosses not explained by *cid* repertoires of females

CI has long been described as a “yes or no” phenomenon in *Cx pipiens*, meaning that crosses are either compatible or fully incompatible [9]. However, after closely investigating the HRs, we recently showed that they could be highly variable in so-called compatible crosses [40]. We further revealed that reduced HRs were associated with cytological signs of canonical CI [40]. We suggested that these phenomena were linked to the *cid* repertoires, as the males infected with *w*Pip-IV, which had distinct *cidB* variants from other *w*Pip groups, induced more intermediate CI than other males.

Studying the cross between Slab females (*w*Pip-III) and Ist males (*w*Pip-IV), we also found intermediate HRs in “compatible” crosses. While on/off compatibility was clearly explained by the presence/absence of specific *cidA* variants, these intermediate HRs did not correlate with any of the *cidA* markers nor with the *Wolbachia* infection rate.

Once again, intermediate HRs were found in crosses involving males infected with *w*Pip-IV. By comparing *cidA* and *cidB* sequences from different *w*Pip groups, we showed that the second interaction interface differed between *cidB* belonging to *w*Pip-IV and *cidB* from other groups, with no variation in the interface II of *cidA* variants (S6 Fig). This could mean that *cidB* variants belonging to *w*Pip group IV never perfectly bind *cidA* variants. The absence of a match at interface II could thus lead to a more unstable *cidA*-*cidB* interaction and explain the intermediate HR values.

### Recombination as the key to the emergence of new CI patterns

A specific *cidB* region, named *cidB-IV-2*, was previously correlated with strong CI-inducing capabilities in males [22,27]. This region is present in the *w*Pip infecting the Ist males studied here. We showed that this specific region is issued from a recombination between the two main groups of *cidB* downstream regions, which are shared by all sequenced *w*Pip groups (Fig 2).

Furthermore, it was previously shown that the only females able to rescue the *modification*s induced by Ist males were those infected with *w*Pip-IV as well as Slab females (*w*Pip-III, studied here) [22,27]. We found that the rescuing ability of Slab females depended on the presence of a specific *cidA* upstream region known as *cidA-beta*: all females with this region can rescue Ist males, while females that have lost it cannot. This variant was also generated by a recombination between two main groups of upstream regions shared by all *w*Pip groups. Here, there were two types of recombinants, the *cidA-beta* and *cidA-IV-delta* regions (Fig 2). When looking at the repertoires found in females infected with *w*Pip-IV, which can all rescue Ist males, we found that they all had one recombinant region in their repertoire (either *cidA-beta* or *cidA-IV-delta*). In other words, all females able to rescue the toxicity of the recombinant region *cidB-IV-2* have a recombinant antidote in their repertoire. Since the *cidA* upstream region (e.g., *beta* or *delta*) interacts with the *cidB* downstream region (e.g., *cidB-IV-2*) in the interaction interface I [21], this finding points to recombinant-for-recombinant binding (Fig 2) in which the recombinant regions can only bind similarly recombinant regions. New CI phenotypes could thus emerge through the recombination of existing variants.

In conclusion, we found that CI evolved in less than 4 years in a *Culex* isofemale line. This evolution is linked to the loss of some *cidA* (antidote) variants in their *cid* repertoire. Our results suggest that these incompatibilities could emerge as a two-step process: first, local changes take place in the CI repertoires while maintaining compatibility with the surrounding mosquitoes, and then migration and secondary contact occur with the incompatible lines. Here, we showed that all repertoires present in a cage were mutually compatible: changes in the repertoires modified the CI pattern only when crossed with an external line. We demonstrated that new compatibility types can emerge rapidly, as a result of the rapid evolution of *Wolbachia* genomes. A common point in the few studies addressing *Wolbachia* microevolution is that both Octomom in *w*Mel [35–37] and *cid* genes in *w*Pip are amplified genomic regions, suggesting the strong role played by gene amplification in the speed of *Wolbachia* genome evolution.

This rapid evolution should therefore be considered when releasing *Wolbachia*-infected mosquitoes, inducing CI, for vector control purposes.

## Materials and methods

All DNA extractions in this study were performed following the cetrimonium bromide (CTAB) protocol [54].

### Isofemale lines

All the mosquito lines used were isofemale lines, i.e., lines created by rearing the offspring resulting from a single egg raft and thus from a single female. We used the following lines: Slab, infected with *Wolbachia* from the phylogenetic group *w*Pip-III, and Istanbul (Ist), infected with *Wolbachia* from the phylogenetic group *w*Pip-IV. Both lines were established in our laboratory in 2005.

We also used six newly established isofemale (sub)lines: two (β+,16+) lines bearing both the *cidA-III-beta* and *cidA-III-16* regions; two (β+,16-) lines bearing only the *cidA-III-beta* marker; and two (β-,16-) lines with neither of these markers. In short, to establish the isofemale line, a cage containing the Slab isofemale line was blood fed. After 5 days, females were isolated in glasses for egg laying. After egg laying, DNA was extracted, and females were genotyped for *cidA-III-beta* and *cidA-III-16* (S5 Table). For each genotype, the two most numerous progenies were kept in order to establish two separate isofemale lines. All isofemale lines were reared in 65 dm screened cages in a single room maintained at 26°C under a 12h light/12h dark cycle. Larvae were fed with a mixture of shrimp powder and rabbit pellets, and adults were fed on a honey solution. Females were fed with turkey blood using a Hemotek membrane feeding system (Discovery Workshops, UK) to enable them to lay eggs.

### Crosses

For each cross performed, around 100 virgin females were put in a cage with around 50 virgin males. After 5 days in cages, females were fed a blood meal; and 5 days later, individual females were isolated for egg laying. This allows us to precisely link a female with its egg raft. Egg rafts were deposited into 24-well plates. The eggs were photographed using a Leica camera and counted using ImageJ. Two days after egg laying, larvae issued from each egg raft were counted manually. HR was calculated as the ratio between the total number of larvae and the total number of eggs. Egg rafts with a null HR were mounted on a slide and examined with a binocular magnifier to ensure that the eggs were fertilized and thus confirm that null HR resulted from CI rather than non-fertilization.

### Repertoire acquisition using Nanopore sequencing of PCR products

We acquired the *cidA* and *cidB* repertoires of eight individuals from the Slab isofemale line, four individuals from the Ist isofemale line, and two individuals from each of the newly established isofemale lines following the pipeline described in [39]. We also acquired *cidA*B combinations for all newly established isofemale lines. DNA was extracted from the individuals. In short, the target sequence was amplified using the adequate primer set (S5 Table, S6 Fig). PCR products were then sequenced by Nanopore sequencing with the MGX Platform (Montpellier) using a MinION (Oxford Nanopore Technologies). Repertoires were then acquired following the pipeline described in [39]. For *cidA*B combinations, the pipeline was adapted: *cidA* and *cidB* repertoires were first obtained, and then the combinations between all *cidA* and *cidB* were artificially created and used as a reference base to run the pipeline.

### *Wolbachia* genotyping

#### Standard PCR

To genotype individuals, DNA was extracted. The presence/absence of both *cidA-III-16* and *cidA-III-beta* was determined using specific PCRs (S5 Table). For individuals negative for both markers, a PK1 PCR (S5 Table) was used to control DNA quality. This was performed on either adult females or fourth instar larvae.

#### Quantitative PCR

Real-time quantitative PCR was run using the LightCycler 480 system (Roche). All DNA samples were analyzed in triplicates for each quantification. A total of 89 females were used for qPCR assays.

The total number of *cidA* copies, along with the number of *cidA-III-beta* and *cidA-III-16* region copies, were quantified using relative quantitative PCR, corrected by wsp, a single-copy *Wolbachia* gene (following [55], primers in S5 Table). Since wsp is present in a single copy per haploid genome, the ratio of the target and wsp signal can be used to estimate the number of target copies per *Wolbachia* genome, correcting for *Wolbachia* infection level and DNA quality. For *cidA-III-beta* qPCR, quantifications were normalized on three distinct individuals with a single *cidA-III-beta* copy.

The infection level (i.e., the number of *Wolbachia* bacteria per female) was assessed with qPCR using a wsp PCR normalized on the *Culex*-specific single-copy ace-2 locus [56]. The ratio between wsp and ace-2 signals can be used to estimate the relative number of *Wolbachia* genomes per *Culex* genome. For *cidA-III-16*/wsp and *cidA*/wsp, the following protocol was used: 3 min at 95°C, 45 cycles of 95°C for 10 sec, 60°C for 20 sec, and 72°C for 20 sec. For *cidA-III-beta/wsp* and *ace2/wsp*, the protocol was as follows: 3 min at 95°C, 45 cycles of 95°C for 3 sec, 62°C for 10 sec, and 72°C for 15 sec. Primer concentrations were 0.6 μM for *cidA-III-16* and *cidA*, 0.4 μM for wsp and ace2, and 1 μM for *cidA-III-beta*.

#### Variant copy number analysis

Variant copy numbers were obtained using both the *cidA* qPCR data and the relative coverage of Nanopore sequencing. Knowing the total number of *cidA* copies, we deduced each variant copy number. For instance, in the case of three *cidA* copies, if out of the two variants present, the coverage of the first was twice the coverage of the second, it could be concluded that there were two copies of the first variant and one copy of the second.

#### Grouping analysis of upstream/downstream regions of variants

Phylogenetic analysis of variants was performed using all the variants already published [22,27,40,41] and present in this study. Network phylogenies of the two regions of interest (*cidA* upstream variable region and *cidB* downstream region, which respectively corresponded to nucleotides 1-800 and 1201-end) were made using Splitstree [57].

#### Nomenclature of *cid* variants

The nomenclature established by [27] was updated, since it could result in identical sequences being given different names in different *w*Pip groups. The fundamentals of the previous nomenclature were retained: names of variants were composed of the name of their upstream region (in Greek letters for *cidA* and in Latin letters for *cidB*) and that of their downstream region (digits). A fixed limit was established between the upstream and downstream regions for both genes (downstream regions starting at nucleotides 801 and 1201 for *cidA* and *cidB*, respectively). A phylogenetic network revealed four clusters of *cidA* upstream regions (Fig 2B), regions within a cluster differing by at most three single-nucleotide polymorphisms (SNPs). We renamed regions so that all those belonging to the same cluster were named using the same Greek letter (e.g., all named alpha). When SNPs were present, to distinguish the sequences, we introduced a number between brackets such as alpha(1). The same was done for *cidB* downstream regions. Similarly, *cidA* downstream regions and *cidB* upstream regions were renamed so that distinct regions had different names, and identical regions had identical names, regardless of the *w*Pip group in which they were found.

For all variant names, the group in which the variant occurs is mentioned. For instance, *cidA-III-alpha(5)-25)* is found in a *w*Pip-III.

The correspondence between the previously used names and the names based on this new nomenclature is provided in S6 Table.

#### Statistical analyses and graphical representation

All statistical analyses were run using R version 4.2.0 [58]. Correlation analyses were performed using a Pearson correlation test. All boxplots and histograms were done using ggplot2 [59], and sequence representation was conducted using the function msaPrettyPrint from the msa package [60].

## Data availability statement

The nucleotide sequences of the new *cidA* and *cidB* variants were deposited in GenBank. Accession numbers are provided in S7 Table. The authors declare that all other data supporting the findings of this study are available in the article and Supporting Information.

## Acknowledgments

We thank Michael Turelli and Nicole Pasteur for their thorough reading and helpful comments on the manuscript and Wei Wang and Mark Hochstrasser for their input on the interface residuals. Sequencing data were generated at the Montpellier GénomiX platform.

## Supporting Information

**S1 Fig. *cidA* and *cidB* variants observed in the Slab isofemale line obtained by Nanopore sequencing**. For each variant, the two variable regions are shown: the upstream region, named with a Greek letter for *cidA* and a Latin letter for *cidB*, and the downstream region, named with a digit for both genes. *cidA* regions, which have a different presence in the repertoires, are represented in green. *cidB* repertoires are common to all repertoires. Females infected with (β+, 16+) or (β+, 16-) *Wolbachia* are able to rescue males infected with *w*Pip-Istanbul (compatible), whereas females infected with (β-, 16-) *Wolbachia* cannot (incompatible).

**S2 Fig. Variations in hatching rates in compatible crosses between Slab females and Istanbul males**. Hatching rates in relation to variations in: A) *Wolbachia* infection level and numbers of copies of B) total *cidA*, C) *cidA-III-beta(2)*, and D) *cidA-III-16*. None of these variations are significantly correlated with hatching rates (Spearman correlation coefficient, *P* = 0.49, 0.28, 0.94, and 0.16 for A, B, C, and D, respectively).

**S3 Fig. Variations of *cidA* copy numbers in *Wolbachia* genomes of Slab females**. A), B), and C) show the numbers of copies of total *cidA*, *cidA-III-16*, and *cidA-III-beta(2)* regions, respectively, in females with repertoires (β-,16-), (β+,16-), and (β+,16+). D), E), and F) show the distribution of *cidA*, *cidA-III-16*, and *cidA-III-beta(2)* copy numbers. All histograms show either bi- or trimodal distributions. All copy numbers were quantified using quantitative PCR relative to the single copy *Wolbachia* gene wsp.

**S4 Fig. Variations of *cidB* copy numbers in *Wolbachia* genomes of Slab females**. *cidB* copy numbers equal *cidA* copy numbers. (β-,16-) *w*Pip has around three *cidB* copies, whereas (β+,16-) and (β+,16+) have around five *cidB* copies. All copy numbers were quantified using quantitative PCR relative to the single copy *Wolbachia* gene wsp.

**S5 Fig. PCR primers used to analyze sequence variation in *cid* genes**. PCRs amplifying the full *cidA* gene (burgundy arrows), variable regions of the *cidB* gene (green arrows), and the *cidA*B tandem (purple arrows) were used for nanopore sequencing of repertoires. PCRs specific to *cidA-III-beta(2)* (yellow) and *cidA-III-16* (orange) were used both in standard PCR and qPCR to determine the presence/absence of these regions.

**S6 Fig. Sequence variation at the *cidA*-*cidB* interaction interface II**. Protein sequences are shown for the span of the interaction interface II (position of the first and last aminoa*cid*s of this interface were obtained thanks to Wang & Hochstrasser, pers. com.). Polymorphic positions are indicated with a gray background. This interface is polymorphic in *cidB*, with *w*Pip-IV *cidB* showing a different sequence from all other groups (sequences shown here are representative of all sequenced variants). By contrast, *cidA* variants are identical in this interaction interface in all sequenced variants regardless of their groups.

**S1 Table. *cid* repertoires of the Istanbul isofemale line**. Nanopore sequencing of *cid* PCR products gave identical repertoires for four distinct Istanbul individuals.

**S2 Table. Crosses between Slab females and Istanbul males**. For each female, we determined the hatching rate (HR) (i.e., ratio between hatched larvae and eggs). The specific *cidA-III-beta(2)* upstream and *cidA-III-16* downstream regions were quantified in each female and normalized over the single copy *Wolbachia* gene wsp. The total *cidA* and *cidB* copy numbers were also quantified and normalized over wsp. Infection rates were assessed by quantifying the single copy gene wsp and normalizing it on the Cx. pipiens single copy gene ace2 (Berticat et al, Proc. R. Soc. Lond. B, 2002).

**S3 Table. Crosses between Slab females from each of the six newly established isofemale sublines (two (β+,16+), two (β+,16-), and two (β-,16-)) and Istanbul males**. The hatching rates and number of hatched larvae are shown for each female. Eggs were only counted in compatible crosses involving (β+,16+) and (β+,16-) females. All females were screened for the presence/absence of two regions of the polymorphic gene *cidA* (*cidA-III-beta(2)* and *cidA-III-16*) to confirm that they had the same repertoire as the female from which the isofemale subline was founded. NF = non-fertilized.

**S4 Table. Reciprocal crosses between (β+,16+) and (β-,16-) isofemale sublines**. Number of eggs and larvae and resulting hatching rates (HR). NF = non fertilized.

**S5 Table. PCR primers and programs used in this study**. qPCR programs are described in the main text.

**S6 Table. Correspondence between variant names in the previous (Bonneau et al, Nat Com 2018) and current nomenclature.**

**S7 Table. GenBank accession numbers of new *cidA* and *cidB* variants used in this study.**

