## Supplementary figures and images for "Intra-lineage microevolution of *Wolbachia* leads to the emergence of new cytoplasmic incompatibility patterns"

### S1 Fig

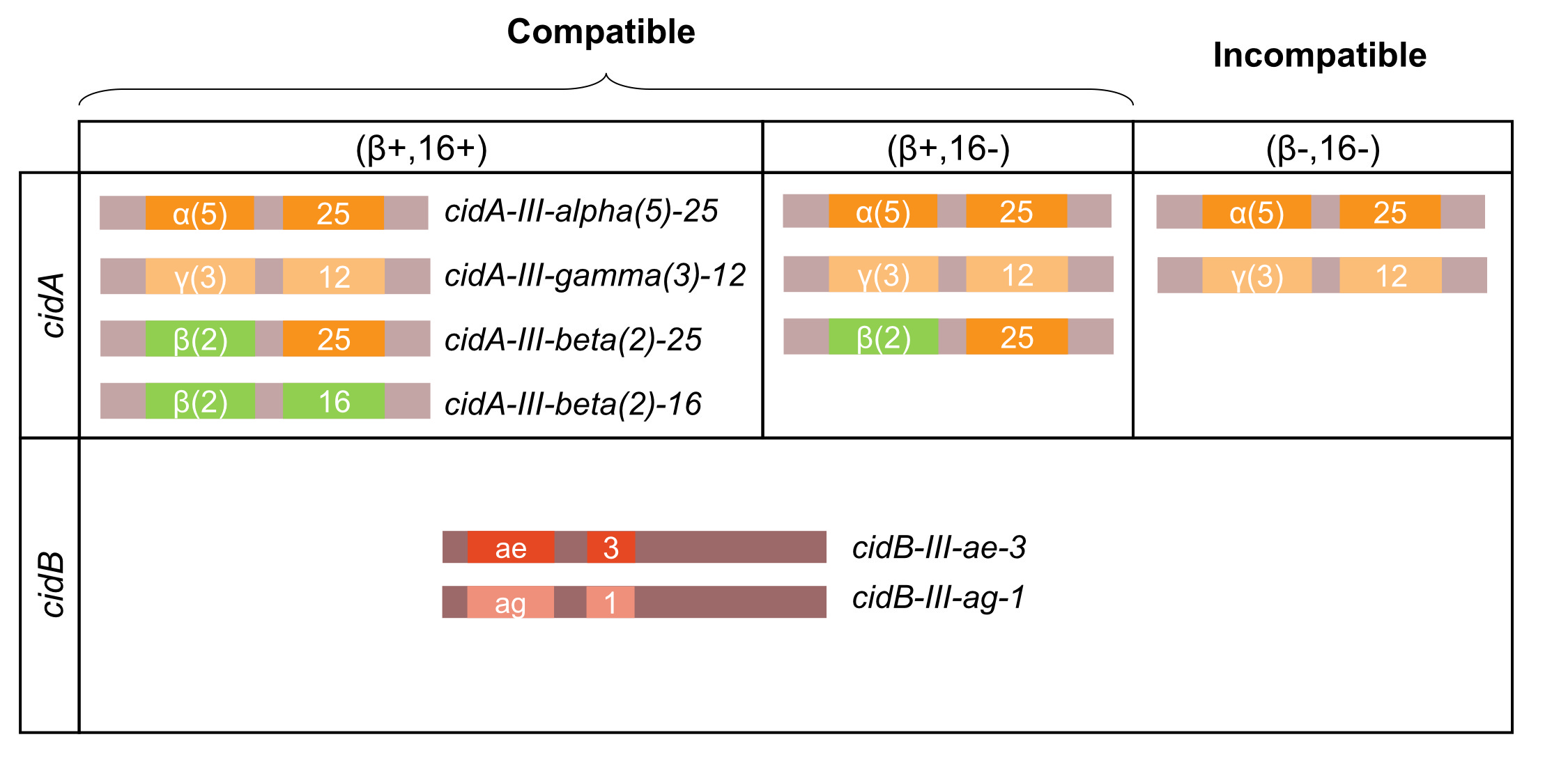

### S2 Fig

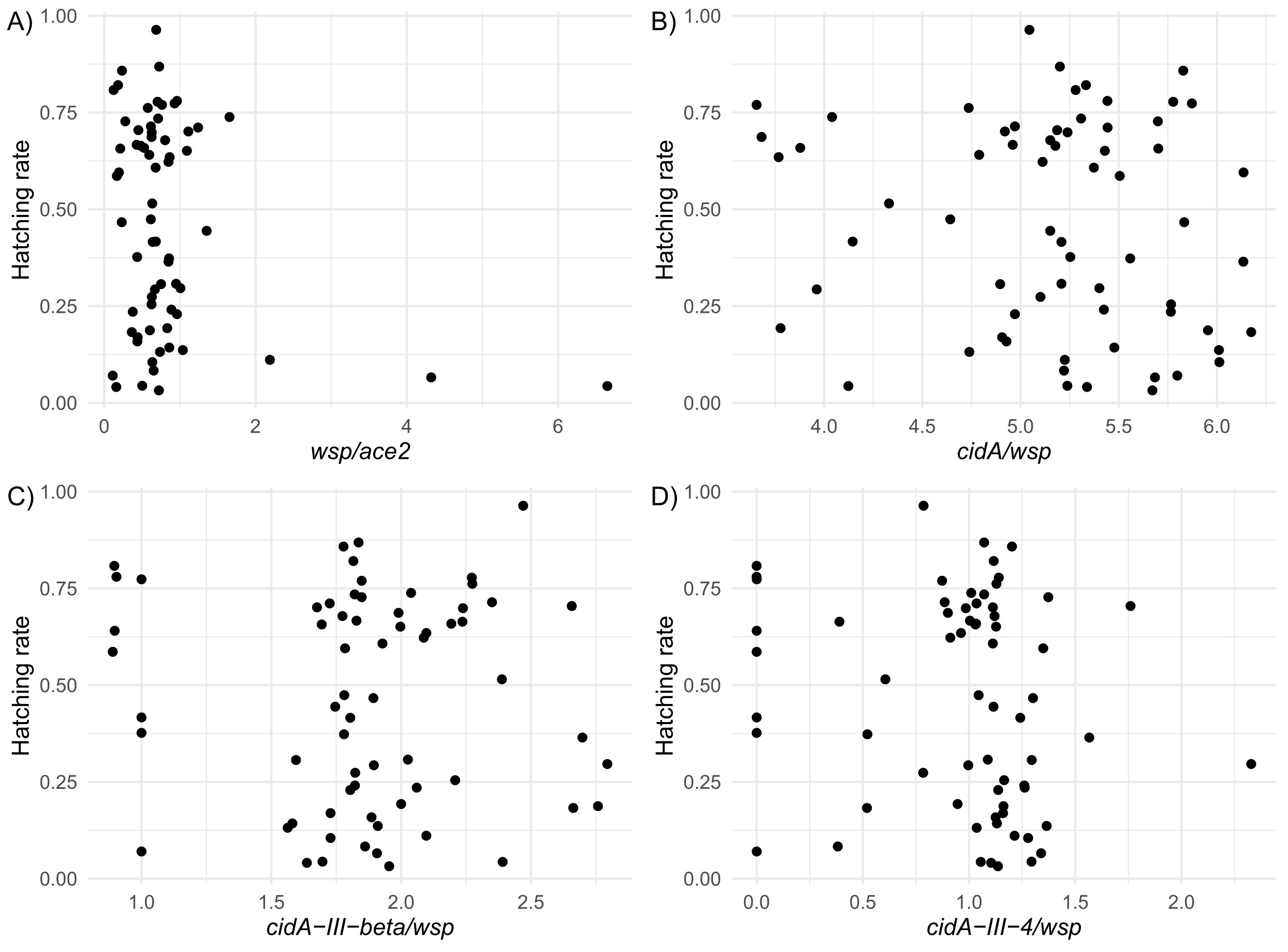

### S3 Fig

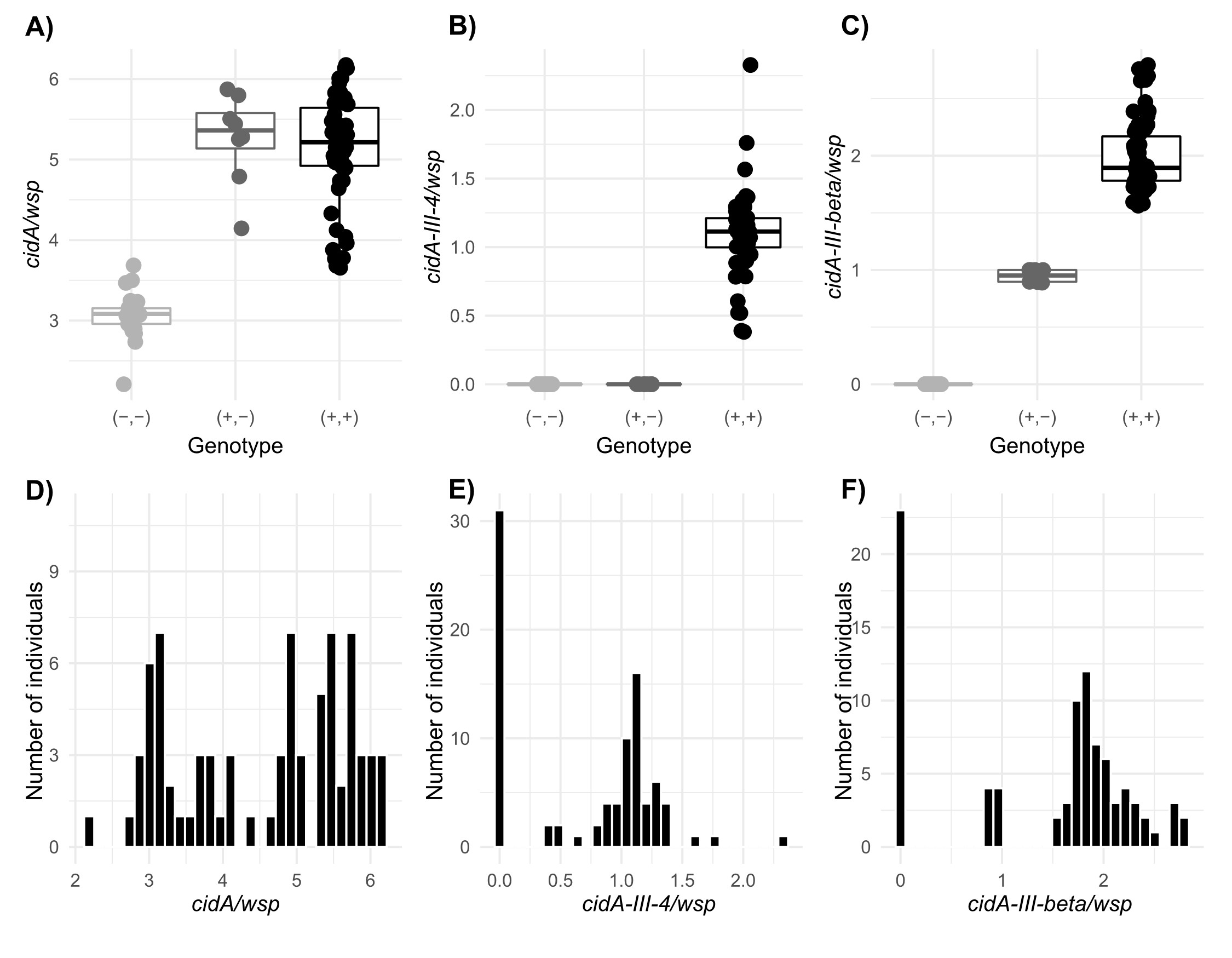

### S4 Fig

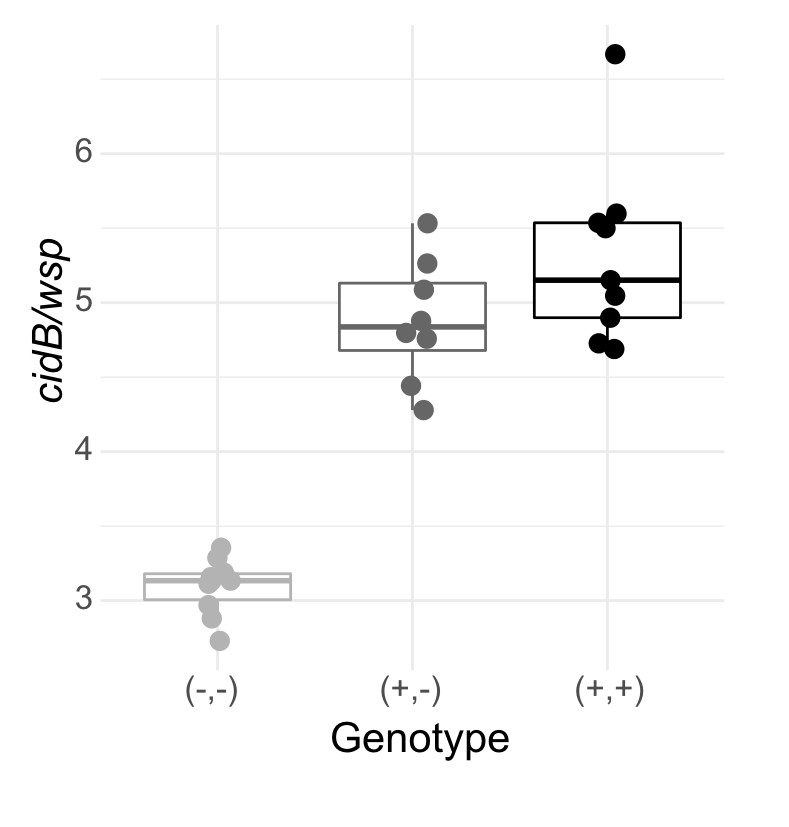

### S5 Fig

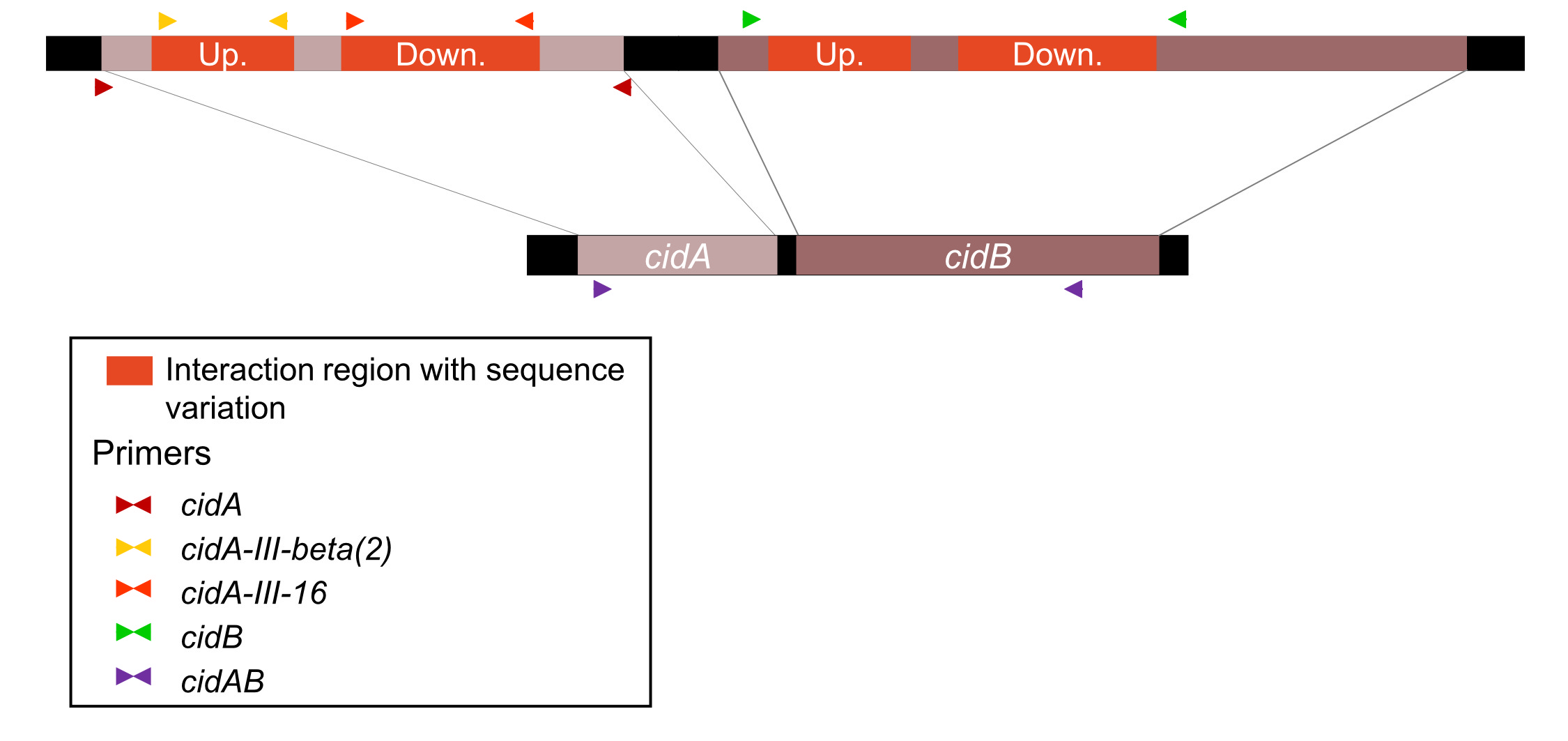

### S6 Fig

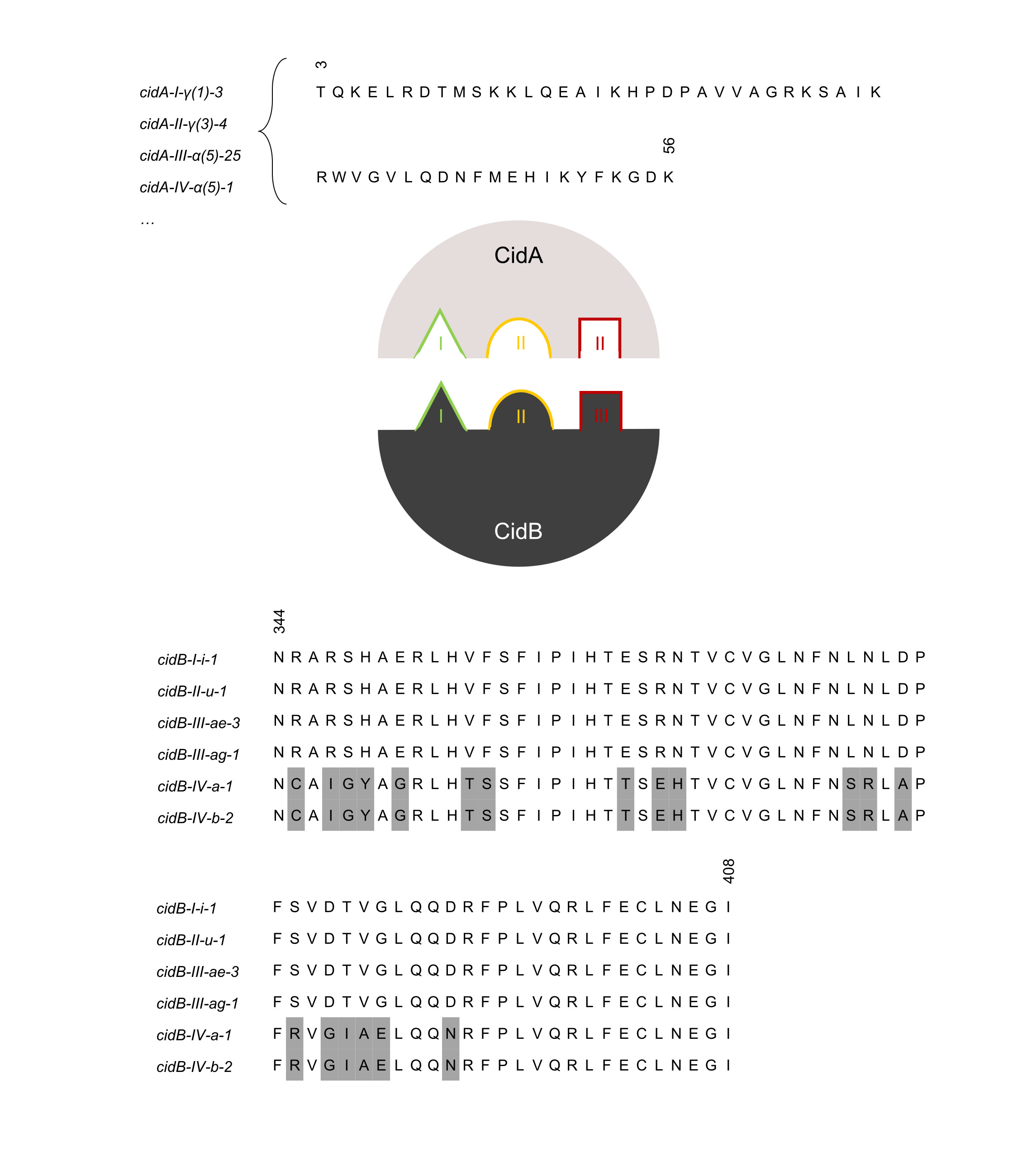
