## Supplementary material for "Intra-lineage microevolution of *Wolbachia* leads to the emergence of new cytoplasmic incompatibility patterns": S1 Table

| ***cidA*** | ***cidB*** |
| --- | --- |
| *cidA-IV-alpha(5)-1*  *cidA-IV-alpha(5)-2*  *cidA-IV-delta(1)-1*  *cidA-IV-delta(1)-2*  *cidA-IV-gamma(5)-1*  *cidA-IV-gamma(5)-2* | *cidB-IV-a1*  *cidB-IV-a2*  *cidB-IV-b1*  *cidB-IV-b2*  *cidB-IV-d1* |
