## Supplementary material for "Intra-lineage microevolution of *Wolbachia* leads to the emergence of new cytoplasmic incompatibility patterns": S5 Table

**Table S5.**

| **PCR** | **Dir primer** | **Rev primer** | **Ref.** | **Tm (°C)** | **Cycles** |
| --- | --- | --- | --- | --- | --- |
| **PK1** | PK1-f  CCACTACATTGCGCTATAGA | PK1-r  ACAGTAGAACTACACTCCTCCA | Duron et al. (2007) | 52 | 33 |
| ***cidA*** | wpip_0282_287981_dir  TGGTCAGGTGTAAGGTTGGA | wpip_0282_289280_rev  CGACCAGAAACACCAAGAGT | Bonneau et al. (2018) | 57 | 31 |
| ***cidB*** | wpip_283_289555_Dir  ACGGCAAACCTAAAAGTTGCT | wpip_283_290818_Rev  TGATGGCTCCTTGTTGTTGC | Bonneau et al. (2018) | 59 | 30 |
| ***cidAB*** | wpip_0282_287994_dir  GGTTGGATAATGCTGCTTTAGGT | wpip_0283_290818_rev  TGATGGCTCCTTGTTGTTGC | This study | 57 | 30 |
| ***cidA-III-beta*** | cidA-IV-gamma_dir1  GATTTGAATCTTATAGGAACAAG | cidA-IV_alpha_rev3  TACAGGACCTTTCCTTCTA | This study | 52 | 30 |
| ***cidA-III-16*** | cidA-I-2-dir1  CTAATTATACAGTTCCTACAAGT | cidA-I-2-rev1  CATAATCTAAGGTCTCTTATTATTAG | This study | 55 | 30 |
| ***cidA* (qPCR)** | wPip282_645_dir2 CGTTCAATCTGTTGAGAAAGA C | wPip282_845_RC2 CCTCCTGCTGAAACTTCTT | This study | 60 | 45 |
| ***wsp* (qPCR)** | Wolpipdir  AGAATTGACGGCATTGAATA | Wolpiprev  CGTCGTTTTTGTTTAGTTGTG | Berticat et al. (2002) | 60 | 45 |
| ***ace2* (qPCR)** | Acequantidir  GCAGCACCAGTCCAAGG | Acequantirev  CTTCACGGCCGTTCAAGTAG | Weill et al. (2000) | 60 | 45 |
| ***cidA-III-beta* (qPCR)** | cidA-IV-gamma_dir1  GATTTGAATCTTATAGGAACAAG | cidA-IV_alpha_rev3  TACAGGACCTTTCCTTCTA | This study | 62 | 45 |
| ***cidA-III-16***  **(qPCR)** | CidA-I-2-dir1000  GCTGCTGTTAATACTTCTAATTA | Cid-A-I-2-rev1161  AAAGCAATCCTTTGGCAAATG | This study | 60 | 45 |
